# Circadian gene-network distortion in high-risk neuroblastoma across multiple biological reference contexts

**DOI:** 10.64898/2026.09.28.754955

**Authors:** N Maussier, D. Tsagkogiannis, N. Baryawno, P. Kogner, J.I. Johnsen, M. Wickström, C. Dyberg, O.C. Bedoya-Reina, T. Sjöberg Bexelius

## Abstract

**Background:** The circadian clock regulates cellular homeostasis, and its disruption has been implicated in aggressive neuroblastoma, particularly in tumours harbouring *MYCN*-amplification. However, it remains unclear whether alterations are restricted to individual clock genes or extend to circadian gene network coordination. We therefore examined circadian clock network disruption in adverse neuroblastoma across multiple biological contexts.

**Methods:** We estimated circadian gene network dysregulation using Delta-CCD in tumours from two neuroblastoma cohorts (SEQC *n*=498 and Kocak *n*=649), comparing clinical features associated with outcome across canonical, adrenal-tissue matched, and developmental references. Robustness was assessed by cross-cohort meta-analysis and leave-one-gene-out analyses. Cox proportional hazards models adjusted for clinical covariates assessed association between individual clock gene expression patient outcome.

**Results:** Delta-CCD was highest in tumours classified as high-risk (study-specific definition) across reference contexts in both cohorts. *MYCN*-amplified tumours showed a more reference-dependent pattern, strongest in adrenal context, while stage 4 tumours showed a similar but weaker pattern. Additional analyses supported the high-risk signal as a distributed network-level alteration rather than a single-gene phenomenon.

**Conclusions:** High-risk neuroblastoma is characterised by robust disruption of coordinated clock gene network organisation across canonical and tissue-matched references, extending beyond individual clock genes. The extent of circadian dysregulation depends on the reference state used.

## Background

Neuroblastoma is a paediatric malignancy of the sympathetic nervous system with marked clinical and biological heterogeneity. While many patients with localised or biologically favourable disease have good survival probabilities, high-risk neuroblastoma remains difficult to cure despite intensive multimodal treatment (1–4). *MYCN* amplification is a hallmark of high-risk neuroblastoma and is strongly associated with adverse clinical outcomes, in part through transcriptional rewiring (3, 4). Recent neuroblastoma studies have emphasised lineage plasticity, adrenergic and undifferentiated mesenchymal-like states, and developmental sympathoadrenal programmes as central features of tumour behaviour (5–9).

The circadian clock-gene network orchestrates rhythmic cellular physiology through coordinated gene expression. This regulation is driven by a cell-autonomous transcriptional-translational feedback loop (TTFL) that generates approximately 24-hour rhythms. Oncogenic signalling can disrupt core circadian clock components, with downstream effects on metabolism, proliferation and therapy response (10–15). The *MYC*-family of transcription factors, including *MYCN* are well-characterised drivers of metabolic reprogramming and circadian disruption in cancer. Experimental evidence from neuroblastoma models demonstrates that *MYCN* activity disrupts the circadian clock by attenuating core clock-gene expression (10, 11). Consistent with this, restoration of the molecular clock is tumour-suppressive in neuroblastoma models (11), and recent work has shown that neuroblastoma cell models differ substantially in circadian rhythmicity and time-of-day drug sensitivity (12). Collectively, these findings suggest that high-risk neuroblastoma may display a perturbation not only of the abundance of individual clock genes, but also of the broader coordination structure of the circadian clock-gene network.

This distinction, between gene expression level and network coordination, is important. A tumour sample collected at an unknown time of day may show high or low expression of a clock gene because of circadian phase, cell composition, transcriptional dysregulation or technical variation. Bulk tumour cohorts rarely include controlled sampling times, making phase-aware circadian inference difficult. Clock correlation distance (CCD) was developed to address this problem by comparing the correlation structure (e.g. in transcript abundance) of clock genes with that of a synchronized reference (13). CCD does not assign a phase to each sample; instead, it asks whether group-level coordination among clock genes resembles or deviates from an expected (e.g. synchronised) reference structure.

Applying CCD to neuroblastoma raises a critical methodological question: which reference should define expected clock-gene coordination? The original CCD framework used synchronised reference data, enabling assessment of deviation from canonical clock progression under controlled conditions (13). This provides a clear circadian-phase reference, but is limited by its cross-species nature and lack of specificity to the developmental and anatomical context of neuroblastoma. Human adrenal tissue offers greater biological relevance because it more closely reflects the tissue of origin of neuroblastoma, although as an unsynchronised tissue, it cannot be interpreted as a pure circadian-phase reference. These strengths and limitations motivated our reference-stratified approach, incorporating a synchronised mouse clock reference as a canonical circadian comparator, human adrenal bulk tissue as a tissue-relevant comparator, and adrenal and developmental references to provide additional context during developmental stages in which circadian oscillatory mechanisms are emerging (7, 16–18).

We therefore set out to investigate whether adverse clinical states in neuroblastoma, namely high-risk disease, *MYCN* amplification and advanced stage, are characterised by distortion of circadian clock-gene network coordination, and whether this signal depends on the biological reference against which it is assessed. To address this, we applied a reference-stratified design across two independent cohorts, using canonical, adrenal and developmental references as complementary comparators.

## Materials and methods

### Study design and reporting framework

This was a retrospective computational study of two publicly available clinical neuroblastoma transcriptomic cohorts (Fig. 1). The primary objective was to test whether adverse clinical groups show altered circadian gene-network coordination relative to favourable comparator groups. Direct Delta-CCD, defined as CCD in the adverse group minus CCD in the comparator group, was used as the main statistical estimate, where positive values indicate greater distance from the reference correlation structure in the adverse group.

**Figure 1.**
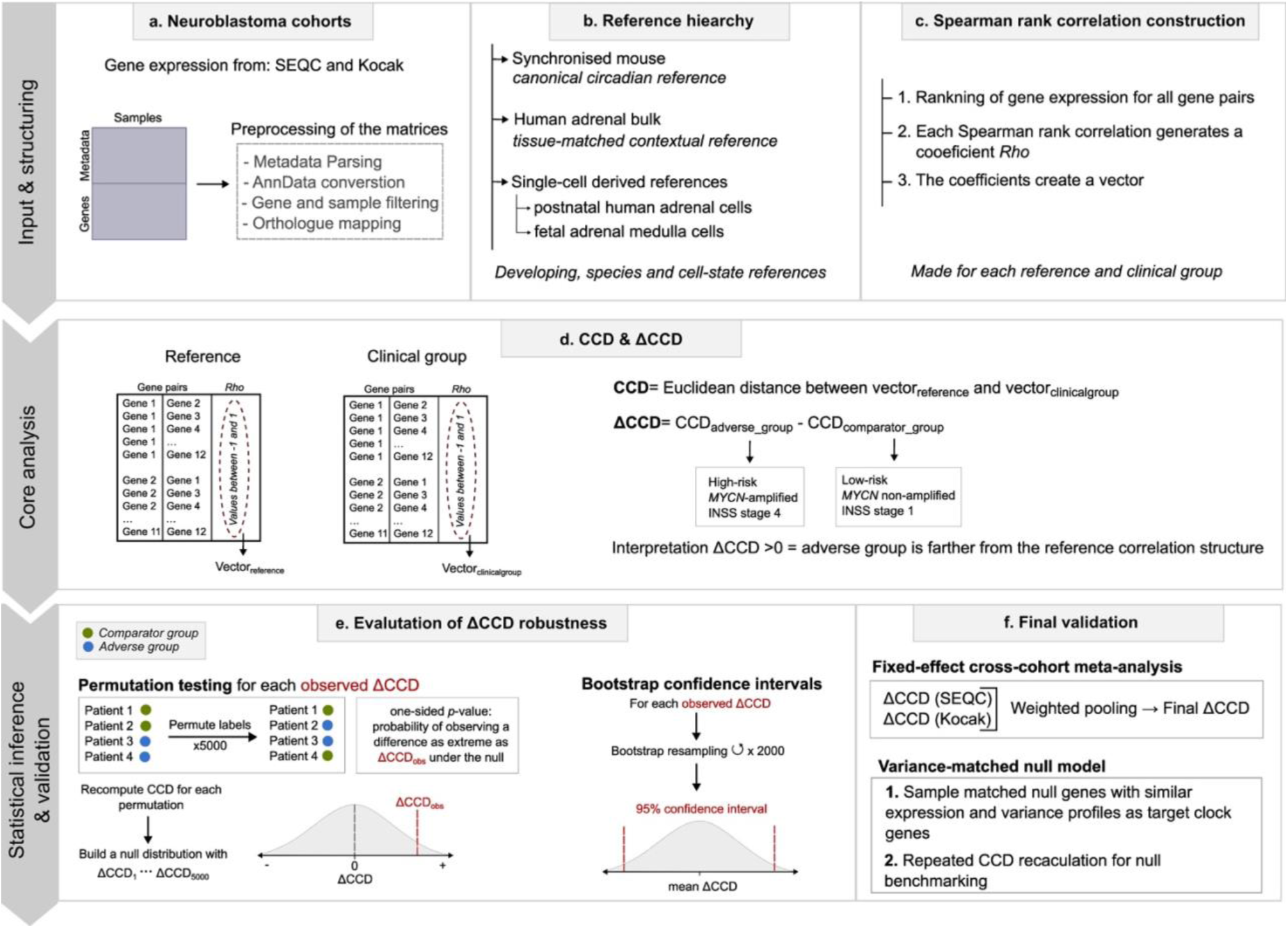
NCC workflow and reference hierarchy. Schematic showing R2-style input matrices, metadata parsing, AnnData conversion, gene and sample filtering, orthologue mapping, Spearman correlation construction, reference correlation meta-analysis, CCD calculation, direct Delta-CCD contrasts, permutation testing, bootstrap confidence intervals, fixed-effect cross-cohort meta-analysis, and unmatched/matched random-gene nulls.

Primary inference focused on the 12-gene circadian panel, two cohorts, log2-transformed expression and direct Delta-CCD testing. The primary clinical contrasts were high-risk versus low-risk, *MYCN*-amplified versus non-amplified and INSS stage 4 versus stage 1. *MYCN*-corrected analyses were conducted using ComBat for *MYCN* amplification status and were only included as technical sensitivity analyses because *MYCN* amplification is part of the biology being studied.

### Neuroblastoma cohorts and clinical variables

The two cohorts were SEQC (GSE62564) and Kocak (GSE45547), accessed through R2 Genomics Analysis and Visualization Platform and associated public repositories (19–22). SEQC contains 498 neuroblastoma tumours profiled by RNA sequencing as part of the Sequencing Quality Control initiative, and Kocak contains 649 tumours profiled using 44K oligonucleotide microarrays from German Neuroblastoma Trials NB90-NB2004. Cohorts were analysed separately before the meta-analysis, to avoid platform-driven artefacts, and concordance across cohorts was used as evidence of robustness.

Clinical variables studied were *MYCN-*amplification status, INSS stage, age-at-diagnosis, sex, vital status and a study-defined harmonised risk variable (Table 1). Because the public datasets did not provide a fully uniform INRG risk classification, a simplified high-risk variable was defined from available information: tumours with *MYCN* amplification or INSS stage 4 were classified as high-risk, and tumours with neither feature were classified as low-risk. This harmonised variable was used for between-cohort comparability and does not replace formal INRG clinical risk stratification. Age was stratified at 18 months where possible. Samples with missing values for a specific variable were excluded only from analyses requiring that variable.

**Table 1.** Clinical characteristics of the neuroblastoma cohorts.

| Characteristics | SEQC (n=498) | KOCAK (n=649) |
| --- | --- | --- |
| **Sex, n (%)** |  |  |
| Male | 287 (58) | 344 (53) |
| Female | 211 (42) | 268 (41) |
| No data | 0 (0) | 37 (6) |
| **Age at diagnosis, n (%)** |  |  |
| < 18 months | 305 (61) | 235 (36) |
| ≥ 18 months | 193 (39) | 414 (64) |
| **MYCN-status, n (%)** |  |  |
| Amplified | 92 (18) | 93 (14) |
| Not amplified | 401 (81) | 550 (85) |
| No data | 5 (1) | 6 (1) |
| **INSS stage, n (%)** |  |  |
| Stage 1 | 121 (24) | 153 (24) |
| Stage 2/2b | 78 (16) | 113 (17) |
| Stage 3 | 63 (13) | 91 (14) |
| Stage 4 | 183 (37) | 214 (33) |
| Stage 4s | 53 (10) | 78 (12) |
| **Risk group, n (%)** |  |  |
| Low-risk | 322 (65) | 407 (63) |
| High-risk | 176 (35) | 242 (37) |
| **Vital status, n (%)** |  |  |
| Alive | 393 (79) | 385 (59) |
| Deceased | 105 (21) | 91 (14) |
| No data | 0 (0) | 173 (27) |

*The harmonised high-risk variable was defined for this analysis and is not equivalent to* *formal INRG risk classification. Abbreviations: INSS – International Neuroblastoma Staging* *System.*

Vital status was considered an outcome-associated grouping rather than a baseline prognostic factor. It was therefore not used as a primary CCD contrast. Survival analyses were performed as contextual analyses of individual genes, univariable and multivariate survival associations were not interpreted as independent prognostic biomarkers.

### Expression processing and gene sets

Expression matrices were analysed in two conditions: a log2-transformed expression, used as the default uncorrected analysis, and *MYCN*-corrected expression. The latter was generated to test whether some coordination patterns persisted after removal of a *MYCN*-amplification-associated expression component; however, it was not interpreted as evidence of *MYCN*-independent biology. The primary circadian gene set comprised 12 core clock-associated genes: *ARNTL*, *CLOCK*, *CRY1*, *CRY2*, *DBP*, *NPAS2*, *NR1D1*, *NR1D2*, *PER1*, *PER2*, *PER3* and *TEF*. This panel was selected to align with the CCD framework and to represent activator, repressor and accessory components of the molecular clock.

### Reference hierarchy and their respective biological interpretation

Two reference classes were emphasised in the main circadian analysis: the synchronised mouse clock reference and the human adrenal bulk reference. The synchronised mouse clock reference was treated as the canonical CCD-compatible circadian reference because it preserves the central feature of the original CCD method: comparison against a synchronized clock-gene correlation structure (13). Its limitation is that it is cross-species and not neuroblastoma-specific.

The non-synchronised human adrenal bulk reference was treated as a tissue-matched contextual reference. Its advantages are species and anatomical relevance, because neuroblastoma arises in the sympathoadrenal developmental axis. Its limitation is that absence of experimental synchronisation means it cannot be interpreted as a direct reference for canonical circadian phase progression. Results against this reference were therefore interpreted as adrenal-context network divergence rather than pure circadian desynchronisation.

Additionally, developmental-state sensitivity analyses were performed using three classes of reference correlation structures derived from published single-cell or single-nucleus datasets: (i) human foetal adrenal medulla references (16), (ii) human postnatal adrenal references (7) and (iii) developing mouse sympathoadrenal lineage references (embryonic stages E9.5-10.5 (17) and E12.5-13.5 (18)). These reference structures were included to assess whether Delta-CCD patterns remained robust across developmental, adrenal and species-specific contexts. Multiple reference-specific Delta-CCD estimates were generated across combinations of cohort, tumour comparison, reference subtype, and expression condition. Consequently, the number of estimates differed between reference classes according to the available reference subtypes and analytical combinations.

The neuroblastoma datasets analysed in this study are bulk transcriptomic cohorts. Single-cell and single-nucleus datasets were used solely to derive reference correlation structures against which bulk tumour samples were evaluated and do not represent *de novo* single-cell profiling of the tumours.

### Computation of CCD

For each reference, cohort, clinical group, expression condition and gene set, pairwise Spearman correlations were calculated across samples for all retained genes (Fig. 1). For the 12-gene circadian panel this yields 66 unique gene-pair correlations. The upper triangle of each correlation matrix was retained and compared with the corresponding reference correlation structure.

CCD was computed as the Euclidean distance between the reference and target correlation matrices after excluding the diagonal and ignoring missing entries. Direct Delta-CCD was then calculated as CCD in the adverse group minus CCD in the comparator group. The adverse groups were high-risk, *MYCN*-amplified and INSS stage 4; comparator groups were low-risk, non-amplified and INSS stage 1. A positive Delta-CCD therefore denotes greater distance from the reference structure in the adverse group.

### Permutation testing, bootstrap confidence intervals and randomisation support

Direct Delta-CCD inference used label permutation and bootstrap resampling within each cohort (Fig. 1). For permutation testing, clinical labels were shuffled while group sizes were retained, group-specific Spearman correlation matrices were recalculated, and a permuted Delta-CCD was recomputed for each draw. The cohort-specific one-sided *p* value tested the pre-specified adverse-greater-than-comparator direction; the cohort-specific two-sided *p* value was retained for transparency.

Bootstrap confidence intervals were estimated by resampling with replacement within each contrasted group, recomputing correlation matrices and Delta-CCD for each bootstrap draw and taking the 2.5th and 97.5th percentiles. The bootstrap standard error was also used in a fixed-effect cross-cohort meta-analysis: cohort Delta-CCD estimates were weighted by inverse bootstrap variance to produce a meta-Delta-CCD, meta *z* statistic and two-sided meta-analysis *p* value.

### Random-gene nulls and leave-one-gene-out robustness

Group-level randomisation analyses were used as a supporting layer rather than as the primary test. Along the lines of the original CCD approach (13), the unmatched random-gene null compared observed group CCD values with CCD values from random gene sets of the same size. The matched-null procedure sampled background genes matched to the target gene set on aggregated mean-expression and variance percentile ranks across target cohorts, with expanding callipers and nearest-rank fallback when needed (Fig. 1).

For the reference-stratified primary table, Benjamini-Hochberg *q* values were calculated from the cross-cohort two-sided meta-analysis *p* values across the main 12-gene tests. Cohort-specific one-sided and two-sided permutation *p* values were retained to show within-cohort support, but the manuscript-level significance statement uses the cross-cohort meta *p* values and meta *q* values.

Sensitivity analyses evaluated direction and significance across reference classes and subtypes (e.g. biological contextual layers); and log2 and *MYCN*-corrected expression conditions. *MYCN*-corrected *MYCN*-status contrasts were flagged as technical sensitivity only because *MYCN*-status was used in the correction step. The leave-one-gene-out sensitivity analyses were used to test whether the primary circadian Delta-CCD signal depended on a single clock gene. For these analyses, one clock gene was omitted at a time; each counted estimate was defined as one expression condition, one cohort, one reference class or subtype, one clinical contrast and one omitted gene. For each analysis, the output retained cohort-specific Delta-CCD, cohort permutation *p* values when present, cross-cohort meta *p* values and BH-adjusted *q* values. The leave-one-gene-out evidence was therefore interpreted as internal robustness of the clock-gene panel rather than as an independent validation cohort.

### Expression and survival context

Clock-gene expression and survival analyses were used to contextualise, not replace, the network-level Delta-CCD results. Cox proportional-hazards models were fitted separately for SEQC and Kocak cohorts using sample-level survival and clinical annotations. Survival variables were resolved before fitting; events were coded as 1 for death/event and 0 for alive/censored. Log2 expression was the primary condition, while *MYCN*-corrected expression was retained only as technical sensitivity. Each clock-gene expression feature was analysed as a continuous predictor and standardised within cohort, so hazard ratios represent the change in hazard per one standard deviation higher expression.

Four Cox specifications were compiled, including univariable models based on expression levels only. Adjusted model A included expression, age group (>=18 months versus <18 months), *MYCN* amplification status and INSS stage covariates. Adjusted model B included expression, harmonised high-risk status and age group. Adjusted model C was an extension of model B that additionally adjusted for sex, but only in cases where the number of survival events was sufficient to support inclusion of the additional covariate. Covariates with zero variance, possible complete separation or high collinearity were dropped. Models were not fitted when event counts or event-per-variable ratios were insufficient. Benjamini-Hochberg *q* values were computed within each condition, cohort, and model. Proportional-hazards assumptions were assessed using rank-transformed Schoenfeld residual tests; both the gene-term *p* value and the minimum *p* value across model terms were retained. Fixed-effect meta-analysis combined SEQC and Kocak gene coefficients by inverse-variance weighting when both cohort-specific models were fitted.

## Results

### Neuroblastoma cohort clinical characteristics

The analysis included 1,147 tumours across the SEQC and Kocak cohorts, comprising 498 tumours from SEQC and 649 from Kocak. *MYCN* amplification was present in 92 of 498 SEQC tumours (18%) and 93 of 649 Kocak tumours (14%), and INSS stage 4 disease in 183 (37%) and 214 (33%) tumours respectively (Table 1). The harmonised high-risk classification identified 176 high-risk tumours in SEQC (35%) and 242 in Kocak (37%). Cohort differences in age distribution and missing vital status were retained rather than imputed.

### Circadian gene-network distortion relative to synchronised mouse clock

Using the mouse canonical circadian reference, high-risk tumours displayed the most pronounced disruption of circadian gene-network coordination. Accordingly, CCD was higher in high-risk than in low-risk tumours across both cohorts: with values of 5.244 versus 4.822 in SEQC, and 5.145 versus 4.854 in Kocak (Fig. 2a, Table 2, Supplementary Tables 1-2). Consistent with this finding, Delta-CCD was positive in both cohorts (SEQC: 0.421, 95% CI 0.198 to 0.642, one-sided permutation *p*=2.00×10^-4^; Kocak: 0.291, 95% CI 0.093 to 0.506, one-sided permutation *p*=0.003; Fig. 2b). The cross-cohort metanalysis remained highly significant (two-sided meta *p*= 5.76×10^-6^, *q*=2.88×10^-5^; Table 2, Supplementary Table 2).

**Figure 2.**
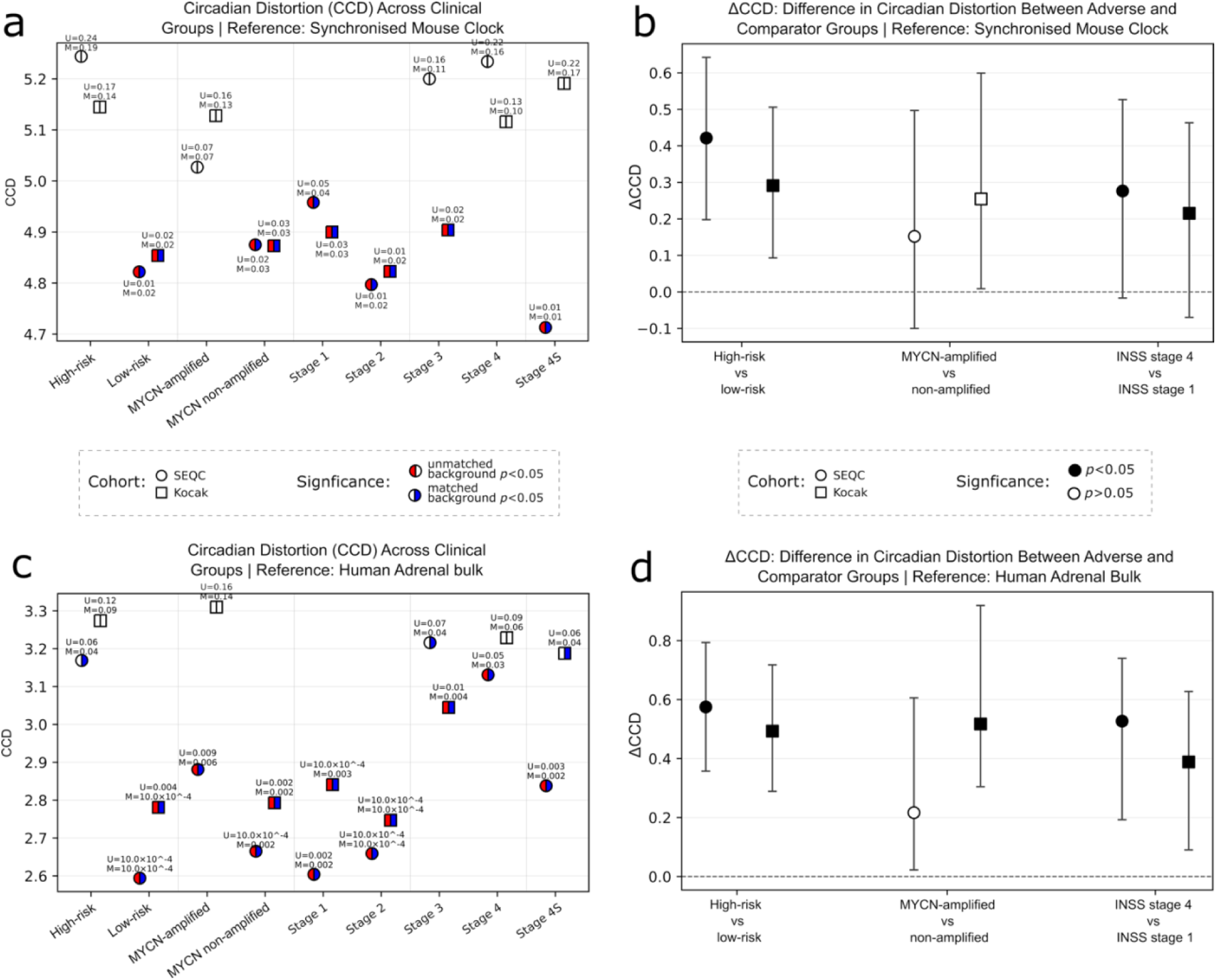
Reference-stratified circadian CCD and Delta-CCD analysis with group-level randomisation support. **a, c)** Group-level circadian distortion (CCD) of the log2-normalised 12-gene circadian panel across clinical groups, using the synchronised mouse clock **(a)** and human adrenal bulk **(c)** as the reference correlation structure. The x-axis shows the clinical groups evaluated against gene-background null distributions, including high risk, low risk, *MYCN*-amplified, *MYCN* non-amplified and INSS stage groups. Each point represents an observed group-level CCD; circles denote the SEQC cohort and squares denote the Kocak cohort. Each point is divided into two halves representing two randomisation tests applied to the same observed CCD: the left half shows the unmatched/random-gene test and the right half the matched-gene test. The two tests use different gene-background null distributions. Red or blue indicates significance at *p*<0.05 for the unmatched or matched test, respectively; white indicates *p*≥0.05. **b, d)** Direct Delta-CCD between the adverse and comparator clinical groups using the synchronised mouse clock reference **(b)** and human adrenal bulk **(d)** as the reference. Delta-CCD is calculated as the adverse group minus the comparator group, with positive values indicating that the adverse group is farther from the reference correlation structure. Points show the observed Delta-CCD, with error bars representing cohort-specific bootstrap 95% confidence intervals. Filled black markers denote cohort-specific one-sided permutation *p*<0.05 for the prespecified adverse-greater-than-comparator direction; open black markers denote *p*≥0.05.

**Table 2.**
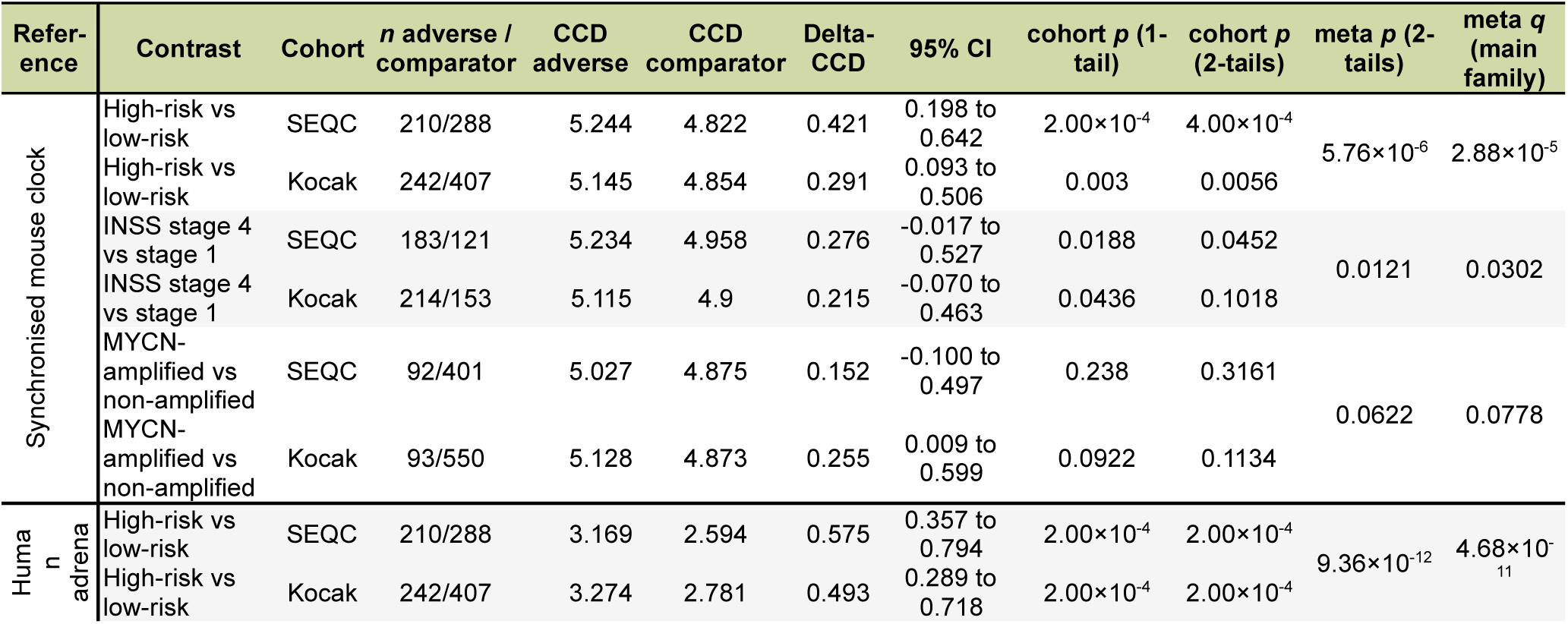
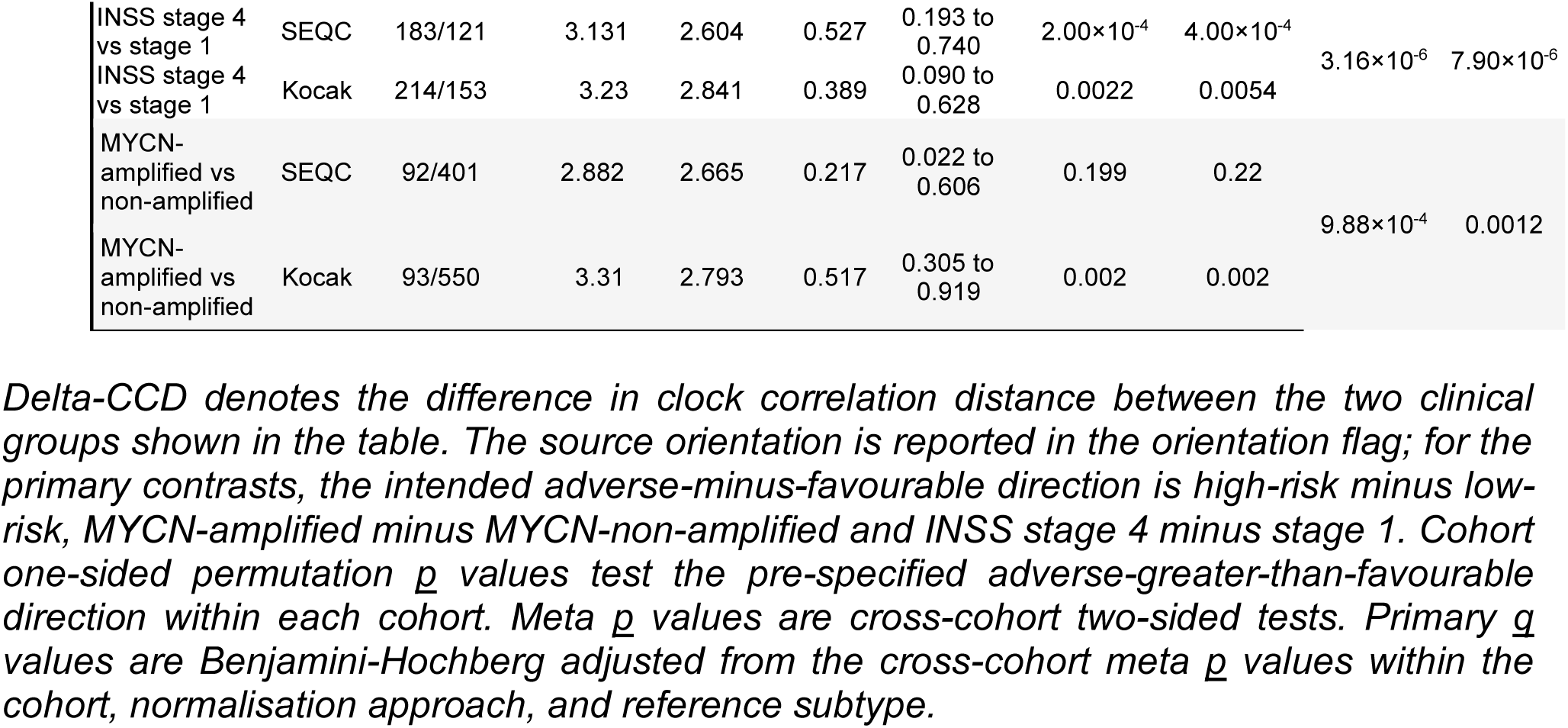
Delta-CCD results stratified by reference dataset for the 12-gene circadian panel (log2 transformed expression).

INSS stage 4 tumours also showed evidence of increased circadian network distortion relative to stage 1 tumours. Delta-CCD was significantly positive in both cohorts (SEQC 0.276, *p*=0.0188; Kocak 0.215, *p*=0.0436), although cohort-specific bootstrap confidence intervals crossed zero (Fig. 2b). Nevertheless, cross-cohort meta-analysis supported a significant association (meta *p*=0.0121, *q*=0.0302; Table 2, Supplementary Table 2), indicating a weaker but reproducible signal compared with the high-risk comparison.

The *MYCN*-amplified tumours showed positive Delta-CCD in both cohorts, but support did not reach the predefined significance threshold (SEQC Delta-CCD 0.152, 95% CI −0.100 to 0.497; Kocak Delta-CCD 0.255, 95% CI 0.009 to 0.599; Fig. 2b; meta *p*=0.0622 and *q*=0.0778; Table 2, Supplementary Table 2). Thus, against the synchronised mouse clock reference, evidence was strongest for the high-risk disease, weaker for INSS stage 4 disease, and not definitive for *MYCN* amplification.

### Circadian gene-network distortion relative to human adrenal bulk reference

The human adrenal bulk reference produced concordant results as observed when using the synchronised mouse clock, but with generally larger effects across both cohorts. Again, high-risk tumours had higher CCD than low-risk tumours (SEQC: 3.169 versus 2.594; Kocak: 3.274 versus 2.781; Fig. 2c, Table 2, Supplementary Tables 1-2) and showed positive Delta-CCD values of 0.575 in SEQC (95% CI 0.357 to 0.794) and 0.493 in Kocak (95% CI 0.289 to 0.718; Fig. 2d), yielding strong cross-cohort support (meta *p*=9.36×10^-12^, *q*=4.68×10^-11^; Table 2, Supplementary Table 2).

For the INSS stage 4 tumours, Delta-CCD was positive in both cohorts (SEQC: 0.527, 95% CI 0.193 to 0.740, one-sided *p*=2.00×10^-4^; Kocak: 0.389, 95% CI 0.090 to 0.628, one-sided *p*=0.0022; Fig. 2d). In contrast to the synchronised mouse reference, cohort-specific bootstrap confidence intervals excluded zero in both cohorts (Fig. 2d), indicating a stronger and more consistent signal of circadian gene-network distortion in stage 4 tumours relative to the human adrenal bulk reference. This association was further supported by cross-cohort meta-analysis (meta *p*=3.16×10^-6^ and *q*=7.90×10^-6^; Table 2, Supplementary Table 2).

In *MYCN*-amplified tumours, Delta-CCD was positive in both cohorts relative to the human adrenal bulk reference (SEQC: 0.217, 95% CI 0.022 to 0.606, one-sided *p*=0.199; Kocak: 0.518, 95% CI 0.305 to 0.919, one-sided *p*=0.002). Unlike the synchronised mouse reference, the cross-cohort association remained significant after multiple-testing correction (meta *p*=9.88×10⁻⁴, *q*=0.0012; Table 2, Supplementary Table 2), indicating stronger support for circadian gene-network distortion in an adrenal tissue context.

### Developmental-context reference analyses

Across developmental-state references, high-risk tumours showed the most consistent evidence of circadian gene-network distortion. Positive Delta-CCD values were observed in 10 of 10 cohort reference estimates, Supplementary Table with significant cross-cohort support in 4 of 5 unique reference contexts (e.g. foetal and postnatal human adrenal gland single-cell-sequenced, human adrenal contextual reference, and full canonical synchronized clock reference; meta *q*<0.05; Fig. 3a, Supplementary Table 2). This pattern was partly retained after *MYCN* correction, with positive Delta-CCD values in 8 of 10 estimates and significant cross-cohort support in 2 of 5 reference contexts. These findings across developmental-stage, tissue-specific, and canonical circadian clock references support circadian network disruption as a robust and context-independent characteristic of high-risk neuroblastoma.

**Figure 3.**
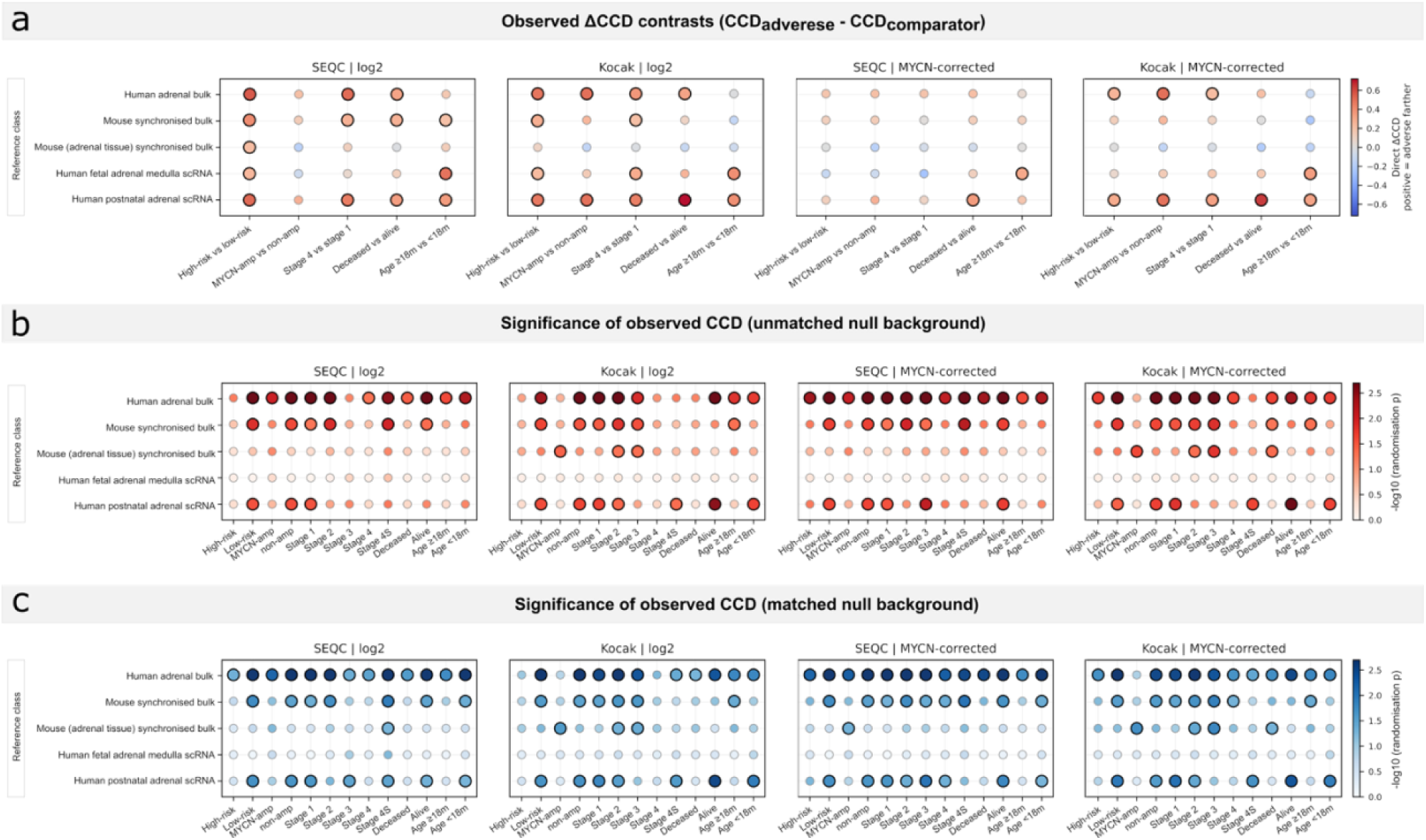
Sensitivity landscape separating direct Delta-CCD and randomisation background support. **a)** Cohort-specific direct Delta-CCD analysis for the 12-gene circadian panel. Columns separate cohort and expression condition: SEQC log2, Kocak log2, SEQC *MYCN*-corrected, and Kocak *MYCN*-corrected. Each point represents one direct adverse-versus-comparator Delta-CCD estimate for a retained reference and clinical contrast. Delta-CCD is calculated as adverse group CCD minus comparator group CCD; red indicates that the adverse clinical group is farther from the reference correlation structure, whereas blue indicates the opposite direction. Larger/bold symbols with black outlines denote statistically supported direct Delta-CCD estimates, defined by cohort-level permutation *p*<0.05. Smaller grey-outlined symbols denote *p*≥0.05. **b)** Significance testing for observed CCD against unmatched gene null background. These panels show group-level randomisation tests, not adverse-versus-comparator Delta-CCD contrasts. Each point represents one clinical group, reference, cohort and expression condition, and tests whether the observed clock-gene CCD for that group is unusual relative to an unmatched random-gene background. Colour intensity represents −log10(*p*). Larger/bold black-outlined symbols denote randomisation *p*<0.05; smaller grey-outlined symbols denote *p*≥0.05. **c)** Significance testing for observed CCD against matched-gene null background. These panels are analogous to panel b but use expression- or gene-property-matched background gene sets where available. Each point represents one clinical group, reference, cohort and expression condition. Colour intensity represents −log10(*p*). Larger/bold black-outlined symbols denote matched-background *p*<0.05; smaller grey-outlined symbols denote p≥0.05. Panels b and c should be interpreted as group-level background support for the CCD signal, not as randomised Delta-CCD contrasts. *Abbreviations: MYCN*-amp – *MYCN*-amplified, non-amp – *MYCN* non-amplified, 18m – 18 months.

Stage 4 versus stage 1 tumours showed a similar but weaker pattern. Positive Delta-CCD was observed in 9 of 10 estimates and significant support in 3 of 5 reference contexts in the log2 analysis, and 7 of 10 positive estimates with support in 2 of 5 reference contexts after *MYCN* correction (Fig. 3a; Supplementary Table 2).

*MYCN*-amplified tumours showed greater variability across developmental references. Positive Delta-CCD was observed 7 of 10 estimates and significant support in 2 of 5 reference contexts in both the log2 and *MYCN*-corrected analyses. The additional clinical contrasts showed intermediate consistency: age ≥18 months versus <18 months was positive in 7 of 10 estimates with support in 2 of 5 reference contexts in both conditions, whereas deceased versus alive status was positive in 8 of 10 estimates with support in 2 of 5 reference contexts in both conditions (Fig 3a; Supplementary Table 2).

### Clock-gene expression and survival outcomes

To evaluate the association between circadian clock gene expression and patient survival, we performed Cox proportional hazards analyses in the SEQC and Kocak cohorts, followed by a meta-analysis combining both datasets. Analyses included univariable models and three multivariable models adjusted for clinically relevant covariates: (A) age group, *MYCN* amplification, and INSS stage; (B) age group and harmonised high-risk status; and (C) age group, harmonised high-risk status and sex.

In the cohort-specific analyses (Fig. 4a, Supplementary Table 3), no significant associations were observed in the univariable analysis of the SEQC cohort, whereas expression of *ARNTL/BMAL1*, *CRY2*, and *PER3* was associated with lower hazard in the Kocak cohort (HR<1, *p*<0.05). After adjustment with multivariable models, only a limited number of genes remained significant. *TEF* showed a protective association in SEQC (HR<1), whereas *CRY1* and *NR1D1* were consistently associated with higher hazard in Kocak (HR>1). *PER2* was associated with increased hazard (HR>1) and *CRY2* with reduced hazard in both cohorts after adjustment for age and risk group (HR<1, Supplementary Table 3).

**Figure 4.**
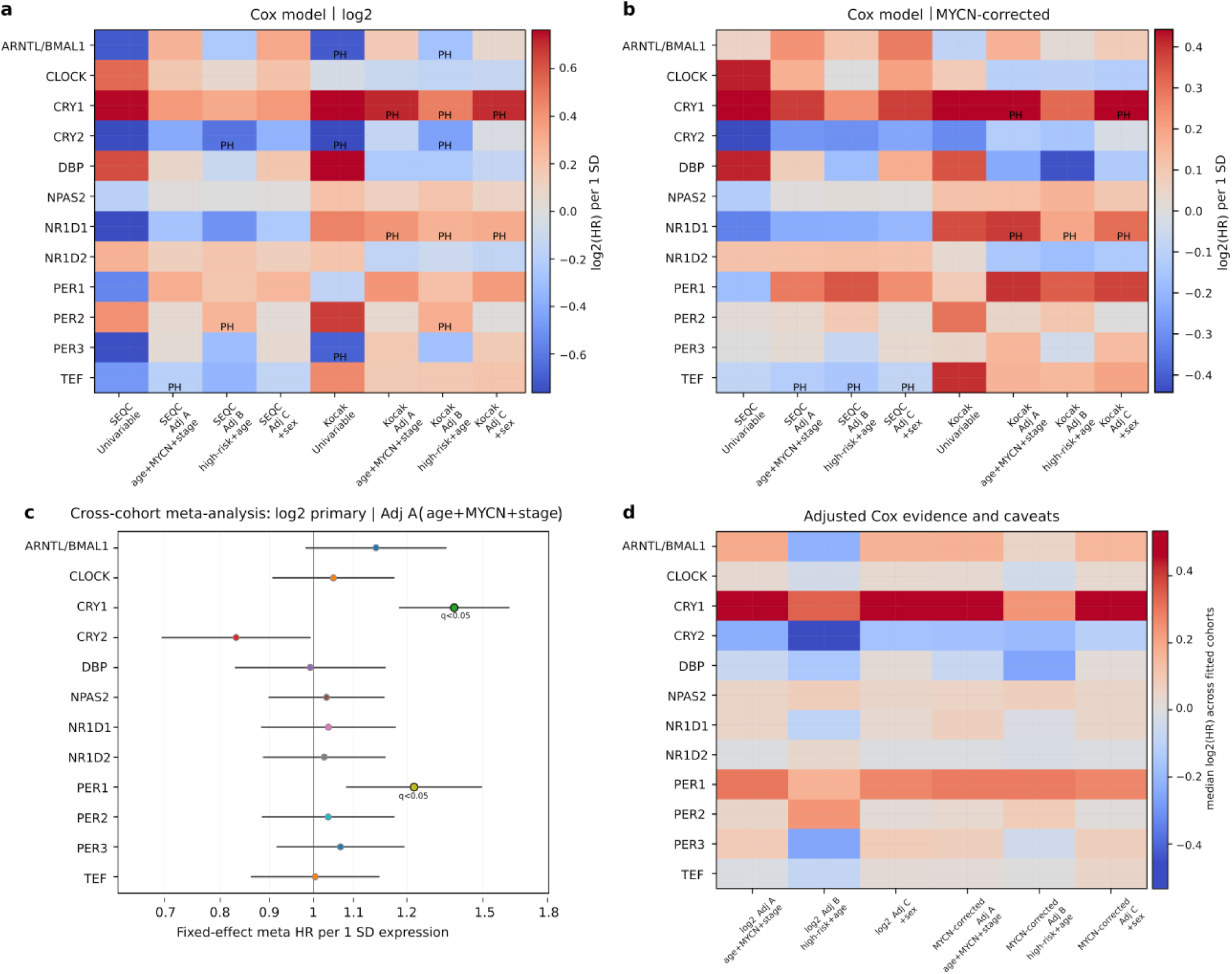
Survival-only clock-gene Cox context and adjusted-model caveats. **a)** Primary log2 Cox survival context for individual circadian-clock genes. Rows show clock genes and columns show cohort-model combinations. Colour indicates the log2 hazard ratio per 1 SD increase in gene expression; red indicates HR>1 and blue indicate HR<1. PH marks gene-term proportional-hazards diagnostic p<0.05, indicating that the hazard-ratio estimate should be interpreted cautiously. Univariable models are shown as survival context only. **b)** *MYCN*-corrected Cox technical-sensitivity context, displayed with the same encoding as panel a. These analyses test whether the direction and approximate strength of gene-survival associations are retained after *MYCN*-status correction, but they should not replace the primary log2 survival analysis. *MYCN*-corrected *MYCN*-related results should be interpreted as technical sensitivity analyses. **c)** Cross-cohort fixed-effect meta-analysis forest plot for the selected expression condition and model, by default log2 primary adjusted model A. Points show fixed-effect meta-analysis hazard ratios per 1 SD gene-expression increase, and horizontal bars show 95% confidence intervals where available. The vertical line marks HR=1. Points to the right of HR=1 indicate higher hazard with higher expression; points to the left indicate lower hazard. Larger black-outlined points denote meta-analysis q<0.05. The default model shown is adjusted model A, defined as expression plus age, *MYCN* status and INSS stage. **d)** Gene-level adjusted Cox evidence and caveats. Rows show clock genes and columns show adjusted model classes across expression conditions. Colour indicates the median adjusted log2(HR) across fitted cohort contrasts.

To assess the contribution of *MYCN*-driven transcriptional effects, we repeated the analyses using *MYCN*-adjusted gene-expression values (Fig. 4b, Supplementary Table 3). Under these conditions, no significant associations were observed in the univariable analyses of either cohort. However, *TEF* remained associated with lower hazard across all adjusted models in SEQC (HR<1), while *NR1D1* and, to a lesser extent, *CRY1* remained associated with higher hazard in the Kocak cohort (HR>1).

To harmonise the assessment of circadian clock gene-survival associations across cohorts and increase statistical power, we subsequently performed a meta-analysis combining the SEQC and Kocak datasets (Supplementary Table 4). In the univariable meta-analysis of the 12-gene circadian clock panel, several genes showed strong associations with survival outcomes. Higher expression of *CRY2*, *ARNTL/BMAL1*, *PER3* and *PER1* was associated with a lower hazard, whereas *CRY1*, *DBP*, *PER2*, *CLOCK* and *NR1D2* were associated with a higher hazard (Supplementary Table 4). Among these, *CRY1* showed the strongest adverse association (HR 1.82, 95% CI 1.60-2.07, *q*=7.3×10^-19^), while *CRY2* showed the strongest protective association (HR 0.45, 95% CI 0.39-0.52, *q*=2.5×10^-24^). *NPAS2*, *NR1D1* and *TEF* were not significant after multiple-test correction (Supplementary Table 4).

Adjustment for clinical covariates attenuated some associations but preserved key signals. In model A (adjusted for age group, *MYCN* amplification, and INSS stage), *CRY1* remained significantly associated with higher hazard (HR 1.40, 95% CI 1.23-1.60, *q*=5.8×10^-6^), while *PER1* showed an adverse association after this adjustment (HR 1.27, 95% CI 1.08–1.50, *q* = 0.021; Fig. 4c). In model B (adjusted for harmonised high-risk status and age group), protective associations persisted for CRY2, *PER3* and *ARNTL/BMAL1*, while CRY1, and PER2 remained associated with higher hazard. Lastly, in model C (adjusted for harmonised high-risk status, age group and sex), *CRY1* remained the only gene consistently significant after within-family correction (HR 1.40, 95% CI 1.22-1.59, *q*=8.0×10^-6^; Supplementary Table 4).

Overall, these results suggest that part of the circadian gene-survival association overlaps with *MYCN*, stage and risk-group structure, but they also support a persistent adverse association for *CRY1* (Fig. 4d; Supplementary Table 3).

## Discussion

Circadian gene-network coordination is fundamentally distorted in high-risk neuroblastoma, extending previous gene-level observations to the level of clock-gene network organisation. This loss of coordination in our study-defined high-risk group was evident against three complementary biological reference frameworks, each answering a distinct biological question. Since high-risk neuroblastoma deviated from all three references, the clock disruption is not aligned with any of the circadian states of any of the biological references used.

These findings raise a broader conceptual question: what constitutes “clock disruption” in this context? Depending on the reference used, deviation may reflect divergence from a canonical synchronised circadian state or from a tissue-matched, developmentally mature or immature state. This distinction is particularly relevant for neuroblastoma as an embryonal malignancy, where a fully established circadian programme may not be present, i.e. an immature circadian null state.

### The biological interpretation of clock disruption depends on reference choice

In our study, delta-CCD functions as a bidirectional variable, further to its original application i.e. measuring both divergence from and similarity to an expected coordinated clock-gene network. The second application enables identification of states resembling distinct tumour characteristics, e.g. answering if low-risk tumours retain a state similar to a developmental state. The interpretation of circadian disruption (delta-CCD) is thus strongly reference-dependent: the synchronised mouse captures deviation from a canonical oscillatory circadian state, while the human adrenal reference offers anatomical relevance to neuroblastoma. Circadian organisation is itself developmentally regulated (23, 24). Therefore, in contexts where rhythmic coordination is immature or absent (23, 24), apparent differences in Delta-CCD could partly reflect underlying developmental timing rather than true clock disruption. In line with this, low-risk tumours having low Delta-CCD compared to both the canonical and adrenal references (Fig. 2c) may indicate that these tumours retain network coordination closer to the reference state, consistent with a more organised, developmentally aligned circadian programme.

Evidence is strongest when the same adverse group shows positive Delta-CCD against both the canonical synchronised reference and the tissue-matched adrenal reference. This criterion was most clearly met by the high-risk comparison. In contrast, *MYCN*-amplified tumours showed stronger support against the human adrenal reference than against the synchronised mouse reference, indicating that reference choice changes whether the result is interpreted as canonical circadian desynchronisation, adrenal-context divergence, or both.

Additionally, our previous findings indicate that MYCN targets associated with poor outcome in neuroblastoma are not homogeneously expressed in all neural-crest-derived cell fates in sympathoadrenal development but rather more likely to be present in neuroblasts and to some extent in Schwann cell precursors, as opposed to chromaffin cells (25). High-risk tumours retained the highest evidence of circadian network distortion, with broadly positive Delta-CCD across human foetal and postnatal adrenal lineages, as well as developing mouse adrenal lineages from the developmental-state reference classes. Stage 4 tumours displayed a similar but weaker pattern, while MYCN-amplified tumours showed greater variability and reduced consistency across developmental references.

### From gene-level associations to network-level disruption

Prior studies have linked *MYCN* with suppression of circadian core transcription factors in neuroblastoma (10, 11) with survival outcomes. In this study, high *CRY1* expression was the only clock-gene expression consistently associated with adverse survival across the multivariable analysis. The variability of these gene-level associations highlights the limitations of interpreting circadian biology through individual transcripts alone and reinforces the value of CCD as a complementary and more robust network-level metric. As circadian function emerges from coordinated feedback loops and phase relationships among core clock genes CCD captures this feature by quantifying distortion of the correlation structure within the clock network rather than focusing on individual gene-expression levels.

The leave-one-gene-out analysis provides an additional robustness check for this network-level interpretation. If the high-risk Delta-CCD result were driven primarily by one clock gene, omitting that gene would collapse or reverse the signal across references. Instead, high-risk disease remained positive across most omitted-gene estimates.

### Limitations

Several limitations require emphasis. CCD is a group-level statistic and is not currently a patient-level biomarker. Bulk tumour expression mixes malignant cells, stromal cells, immune cells, vascular components and differentiating compartments. The single-cell-derived references reduce neither the need for direct tumour single-cell analysis nor the risk that reference correlations reflect developmental or cell-state gradients rather than circadian phase. Survival analyses are more mature after the corrected Cox compilation, because fitted adjusted models and cross-cohort meta-analysis are now available. Nevertheless, proportional-hazards diagnostics and adjustment-dependent gene effects mean that the Cox results should be presented as supportive clinical context rather than as a standalone prognostic signature.

## Conclusions

In conclusion, this study demonstrates that high-risk neuroblastoma is associated with reproducible distortion of circadian gene-network coordination across two independent cohorts. The high-risk signal was robust across canonical circadian, tissue-contextual and developmental reference frameworks(whereas INSS stage 4 disease showed a concordant but weaker pattern and *MYCN*-amplified tumours displayed greater reference dependence)The biological distinct reference-stratified design highlights that circadian disruption is not a single biological entity and our findings emphasize that neuroblastoma may occupy distinct circadian states ranging from canonical desynchronisation to developmental immaturity or absence of a fully established circadian network. Rather than assuming a single disrupted clock phenotype, it may be more appropriate to consider whether distinct tumour-specific circadian states exist, including states with minimal or absent rhythmic organisation. Defining such phenotypes could be critical for translational strategies: if some tumours reflect a circadian “null” state, approaches based on re-synchronisation may be ineffective, and alternative strategies targeting more fundamental aspects of clock biology or developmental state may be required (12).

Future studies should determine whether *MYCN* directly perturbs circadian network coordination in synchronised neuroblastoma models and further define tumour-specific circadian states before these findings can be translated into therapeutic strategies.

## Acknowledgements

The authors thank the investigators who generated and shared the SEQC, Kocak and adrenal/developmental reference datasets and the developers of the R2 platform. Part of the computations and data handling were enabled by resources provided by the National Academic Infrastructure for Supercomputing in Sweden (NAISS), partially funded by the Swedish Research Council through grant agreement no. 2022-06725.

## Additional information

### Authors’ contributions

Conceptualisation, O.C.B.-R., T.S.B.; methodology, O.C.B.-R., T.S.B., N.M.; software, O.C.B.-R., N.M.; formal analysis, N.M.; investigation and data curation, N.M., O.C.B.-R; writing – original manuscript, N.M.; visualisation, N.M., O.C.B.-R.; writing – review and editing, O.C.B.-R., T.S.B., N.M., D.T., N.B., P.K., J.I.J., M.W., C.D.; supervision, O.C.B.-R., T.S.B.; funding acquisition, T.S.B.

### Ethics approval and consent to participate

This study used publicly available, anonymised gene-expression and clinical metadata. No new human participants, human tissue or identifiable individual data were collected.

### Consent for publication

Not applicable; the manuscript does not contain identifiable individual participant data.

### Data availability

The primary datasets analysed in this study are available through GEO/R2, including GSE62564 and GSE45547. Analysis scripts, manifests and derived result tables are deposited in the public GitHub repository https://github.com/BobsYourOnco/Maussier_et_al_2026.

### Competing interests

The authors declare no competing interests.

### Funding information

This work was supported by grants from the Swedish Childhood Cancer Foundation (PR2021-0129 to O.C.B.-R.); KI Funds (2022-01925 to O.C.B.-R.). O.C.B.-R. was supported by an Assistant Professorship from the Swedish Childhood Cancer Foundation (TJ2021-0137).

### Use of artificial intelligence tools

Large language model was used to revise the grammar of the manuscript. The authors are responsible for the content.

## Supplementary information

### Supplementary methods

#### Nina’s Circadian Clockwork (NCC) workflow, CCD and Delta-CCD implementation

Input expression files were handled using the Nina’s Circadian Clockwork (NCC) workflow, which reads R2-style expression matrices, separates sample-metadata records from gene-expression records, converts the data into AnnData objects and applies cohort-level sample inclusion or exclusion filters before correlation analysis. Genes were mapped between reference and target species using a one-to-one orthologue table where needed. This table was obtained from the ENSEMBL repository version 116 using high-confidence 1:1 orthologue between human (GRCh38.p14) and mouse (GRCm39). Mouse and mouse-derived reference correlation matrices were constructed using reference-species gene symbols. In detail, human genes were mapped to human 1:1 orthologue before target correlations were calculated; therefore, mouse *Bmal1* corresponds to human *ARNTL* in the manuscript analyses. Strict gene matching was used for the core analyses, and analyses were restricted to genes present in both the reference and target matrices.

#### Software, compilation and reporting units

Analyses were implemented using Python workflows, including the Nina’s Circadian Clockwork (NCC) Delta-CCD and matched-null workflow. The NCC workflow generated reference correlations, target correlations, direct Delta-CCD contrasts, permutation draws, bootstrap confidence intervals, fixed-effect meta-analysis statistics, unmatched random-gene nulls and matched-null summaries. Compiled tables retained log2 and *MYCN*-corrected conditions only.

For direct Delta-CCD estimates, the present article reports cohort-specific Delta-CCD, sample counts, adverse-group CCD, comparator-group CCD, one-sided and two-sided permutation *p* values, bootstrap confidence intervals, cross-cohort two-sided meta-analysis *p* values and BH-adjusted meta *q* values. Repeated meta *p* and *q* values across SEQC and Kocak table entries are expected because the same cross-cohort test is displayed next to the cohort-specific effect sizes.

**Supplementary Table 1.** log2-normalised and *MYCN*-corrected CCDs with references, cohorts, clinical variables, and randomisation parameters and support for variance-matched and unmatched gene expression.

**Supplementary Table 2.** log2-normalised and *MYCN*-corrected Delta-CCDs with references, cohorts, contrasts, *p* values, *q* values, and meta-analysis results.

**Supplementary Table 3.** Clock-gene Cox survival context, including model specifications, fit status, hazard ratios, confidence intervals, *p* values, *q* values, proportional-hazards diagnostics.

**Supplementary Table 4.** Clock-gene Cox survival fixed-effect SEQC/Kocak meta-analysis, including model specifications, fit status, hazard ratios, confidence intervals, *p* values, and *q* values.

**Supplementary Figure 1.**
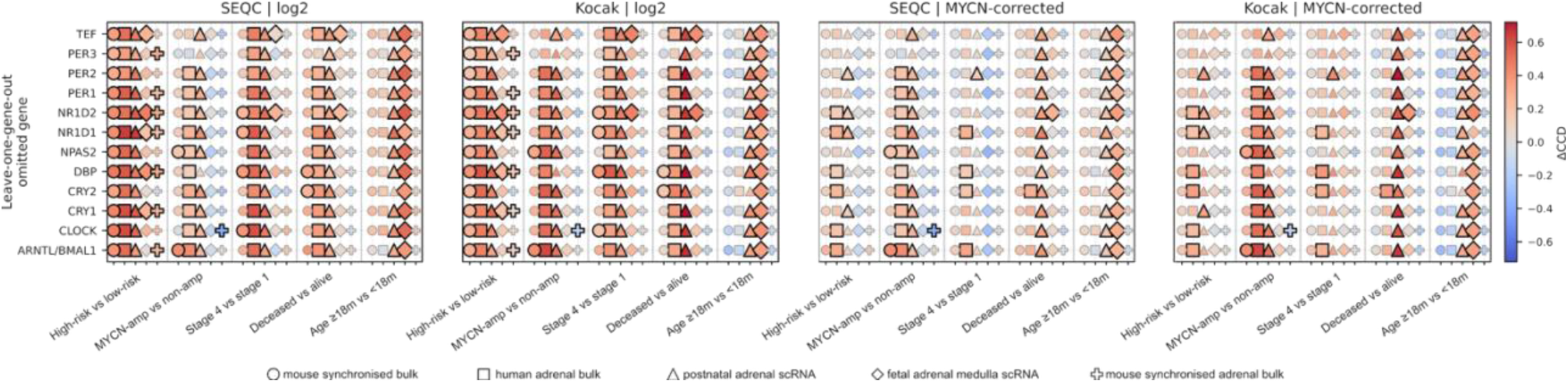
Leave-one-gene-out robustness analysis. Each panel uses the same cohort-by-condition layout as Figure 3 panels a-c. The x-axis is organised by clinical contrast, with separate internal columns for each retained reference; the y-axis indicates the omitted clock gene. Each point represents one contrast-reference-omitted- gene leave-one-out Delta-CCD estimate. Colour shows the signed leave-one-gene-out Delta-CCD after omitting the indicated gene. Red indicates that the adverse group remains farther from the reference correlation structure after omission of that gene; blue indicates attenuation or reversal. Larger/bold black-outlined symbols denote statistically supported leave-one-gene-out estimates; smaller grey-outlined symbols denote non-supported estimates. Human *ARNTL* and mouse *Bmal1/Arntl* are harmonised as *ARNTL/BMAL1*. Symbol shape denotes the retained reference used for the estimate: circles represent the mouse synchronized bulk reference; squares represent the mouse synchronized adrenal bulk reference; upward triangles represent the human adrenal bulk contextual reference; diamonds represent the postnatal human adrenal single-cell reference; and filled pentagons represent the foetal adrenal medulla single-cell reference. *MYCN*-corrected *MYCN*-status comparisons are shown only as technical sensitivity analyses because *MYCN* status was used during the correction step.

## Notes

### Competing Interest Statement

The authors have declared no competing interest.

