## Supplementary figures and images for "Circadian gene-network distortion in high-risk neuroblastoma across multiple biological reference contexts"

### Supplementary Figure 1

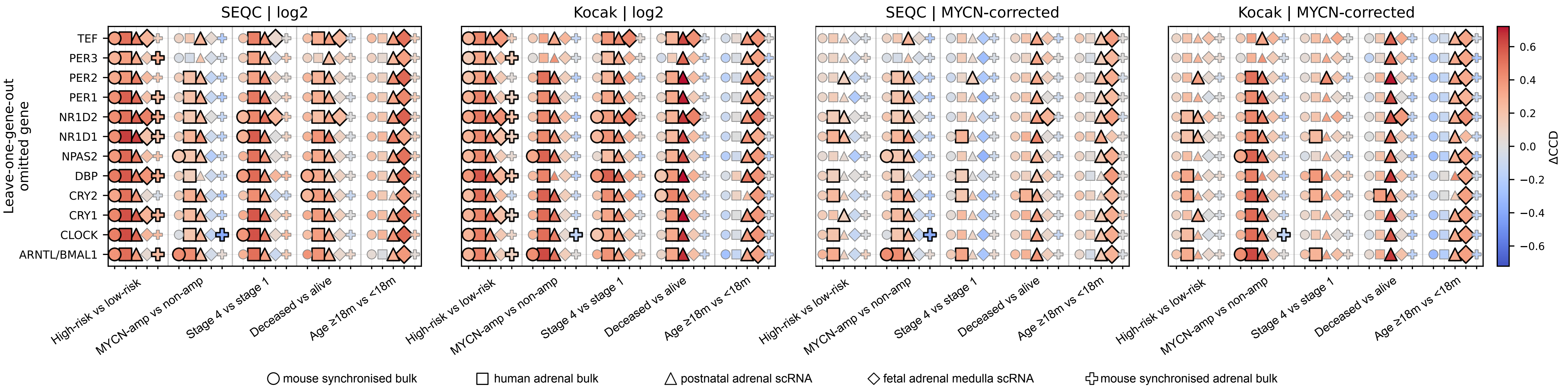
